# BxbI-mediated insertion of a 77kb human RET sensitive haplotype into the mouse genome to generate a humanized model of Hirschsprung disease

**DOI:** 10.64898/2026.07.05.736620

**Authors:** Ryan D. Fine, Benjamin E. Low, Jarod Rollins, Jon M. Laurent, Michael V. Wiles, Aamir Zuberi, Jef D. Boeke, Aravinda Chakravarti

## Abstract

Hirschsprung disease (HSCR) is a complex developmental disorder of the enteric nervous system, primarily driven by regulatory variants within enhancer elements of the *RET* gene. To investigate how these variants lead to aganglionosis, we developed a humanized mouse model by inserting an intact 77kb human *RET* genomic locus into the *Rosa26* safe-harbor locus. Utilizing “big DNA” synthetic biology and Bxb1-mediated recombination, we integrated the complete human locus including all exons, introns, and upstream regulatory elements which we validated via nanopore and short-read sequencing. Functional analysis confirmed *in vivo* human *RET* expression; however, our initial HSCR-associated “sensitive” haplotype expressed at only 21% of wild-type levels. This significant reduction proved insufficient to rescue the viability when endogenous mouse *Ret* was deleted. We identified that this deficiency is partially driven by five risk SNPs within established enhancers. Specifically, using CRISPR/Cas9 to restore a conserved So×10 binding site (converting a sensitive SNP to a protective one) increased *RET* expression by 1.9-fold and restored transcription factor binding. This study provides a robust framework for modeling human-specific regulatory disorders and demonstrates the critical impact of non-coding variation on disease pathogenesis.

## Introduction

One of the major and long-standing questions in all genetics has been to understand how a genotype gets elaborated into a phenotype. In contrast to simple Mendelian traits where every genotype leads to a specific phenotype, the path/s from multi-locus genotype to a polygenic or multifactorial (complex) trait is complex and often “inconstant and modifiable.”^1^ The first genetic dissection of such a complex trait was *truncate wing* in Drosophila melanogaster by Altenburg and Muller (1919)^1^ who used precisely these descriptors to describe how one genotype could lead to different phenotypes.

The recent successes of genome-wide association studies to map the locations of thousands of traits^2^ reveals intense polygenicity and has refocused our attention on this problem. As originally postulated by Fisher^3^ any genotype can produce widely distinct phenotypes while any phenotype (class) is composed of many distinct genotypes. In other words, individuals, and their molecular machinery, can distinguish genomes with say 5 disease-risk increasing variants from those with 50 even though neither produce a deterministic outcome.^4^

In this study, we focus on Hirschsprung disease (HSCR) as an exemplar disorder to begin to understand this complex problem. HSCR is a neonatal developmental disorder of the enteric nervous system (ENS) that exhibits a large range of genotypic and phenotypic diversity.^5,6^ The disorder is clinically diagnosed by an absence of intestinal neuronal ganglia in the surrounding submucosal plexuses, primarily in the distal colon, however, affected individuals, as well as first degree relatives, show significant variation in the extent of aganglionosis from short segment (S-HSCR) absence in the sigmoid colon to long segment (L-HSCR) absence up to the splenic flexure or even total colonic aganglionosis (TCA).^5,6^ Overlaid on this variable extent of aganglionosis, many individuals with pathogenic variants in HSCR sensitive genes^7^ exhibit incomplete disease penetrance within families.^8^

Fortunately, the molecular basis of HSCR pathogenesis is fairly well understood with a minimum of 24 genes and 9 loci explaining 72% of its population attributable risk.^7,9^ Remarkably, 67.2% of this risk is mediated through a single gene regulatory network (GRN) that regulates the expression of *RET* (a receptor tyrosine kinase gene) and *EDNRB* (a G-protein coupled receptor gene) genes that are rate-limiting for ENS development.^10,11^ Within this GRN, *RET* plays the most significant role because beyond the risk it imparts from pathogenic coding variants its largest risk contributions to HSCR is from multiple, common, hypomorphic variants that disrupt the activities of transcriptional enhancers of *RET* expression in the developing gut.^12^ As is hypothesized for other complex disorders, HSCR risk is largely from multiple hypomorphic regulatory variants. In this study, our goal was to understand the molecular processes that convert these regulatory genotypes into disease by affecting the cascade of regulatory genotypes affecting chromatin, affecting gene expression, followed by changing protein expression, inducing dysregulation of the GRN and affecting differentiation and proliferation of enteric neural crest cells destined to populate the gut.

In this study, we successfully engineered a mouse strain containing a 77kb human *RET* sensitive haplotype with its entire suite of introns, exons, and enhancers. Our aim was to dissect the contributions of *RET* regulatory variants to gene expression and the ENS phenotype. Furthermore, in an attempt to expose and test this genes activity we also deleted the mouse endogenous *Ret* gene making this strain dependent on human *RET* function alone. Surprisingly, despite detecting transcription and translation of human *RET* in the expected tissues, the 77kb humanized allele bearing a fully intact promoter and wildtype coding sequence is not sufficient to rescue the loss of the endogenous mouse *Ret* locus. This is in part due to presence of 5 sensitive/risk SNPs located within enhancer sites as mutation of a single SNP is sufficient to induce a 5% increase in *RET* expression and increase binding of the known *RET* transcription factor, So×10. Collectively, our results suggest that simple copy/paste of human genes in an exogenous system is not sufficient to fully recapitulate endogenous gene expression. We hypothesize this may be due to lack of additional enhancer elements, human transcription factors (TFs) or even RET’s co-receptor and ligand, GFRA1 and GDNF.

## Results

To design the human *RET* locus, we built upon our primary knowledge of previous work that had identified 10 of 38 GWAS-implicated variants with differential enhancer activity in luciferase plasmid expression assays in human neuroblastoma SK-N-SH cells.^12^ This led to our first choice of a *RET* locus design of 179kb that extended 500bp beyond the 3’ UTR and 500bp beyond a differentially accessible RARB binding disease variant, rs2506030 (**Figure 1A**). This locus would incorporate 7 of the 10 known HSCR-associated and differentially accessible enhancer SNPs as well as all 5 enhancers known to bind the 4 known developmental ENS transcription factors (TFs), RARB, GATA2, SOX10, and PAX3.^10–12^ The risk-increasing alleles at the RARB, GATA2, and SOX10 binding sites collectively have an odds ratio of 4.4 (p=3.62×10^-22^).^10^ Despite the clear importance of all three of these enhancers, only the SOX10 site, rs2435357, exhibits sequence conservation across species (**Figure 1A**). This provided us the motivation to build a fully intact human locus in mice. We also designed a smaller 77kb locus incorporating 5 of these variants bearing just the GATA2, SOX10, and PAX3 sites with a full suite of HSCR-risk alleles at all 5 variants with a corresponding odds ratio of 3.1 (P=8.31×10^-10^).^10^ We herein refer to this 77kb locus as *RET-*sensitive (*RET-s*).

**Figure 1.**
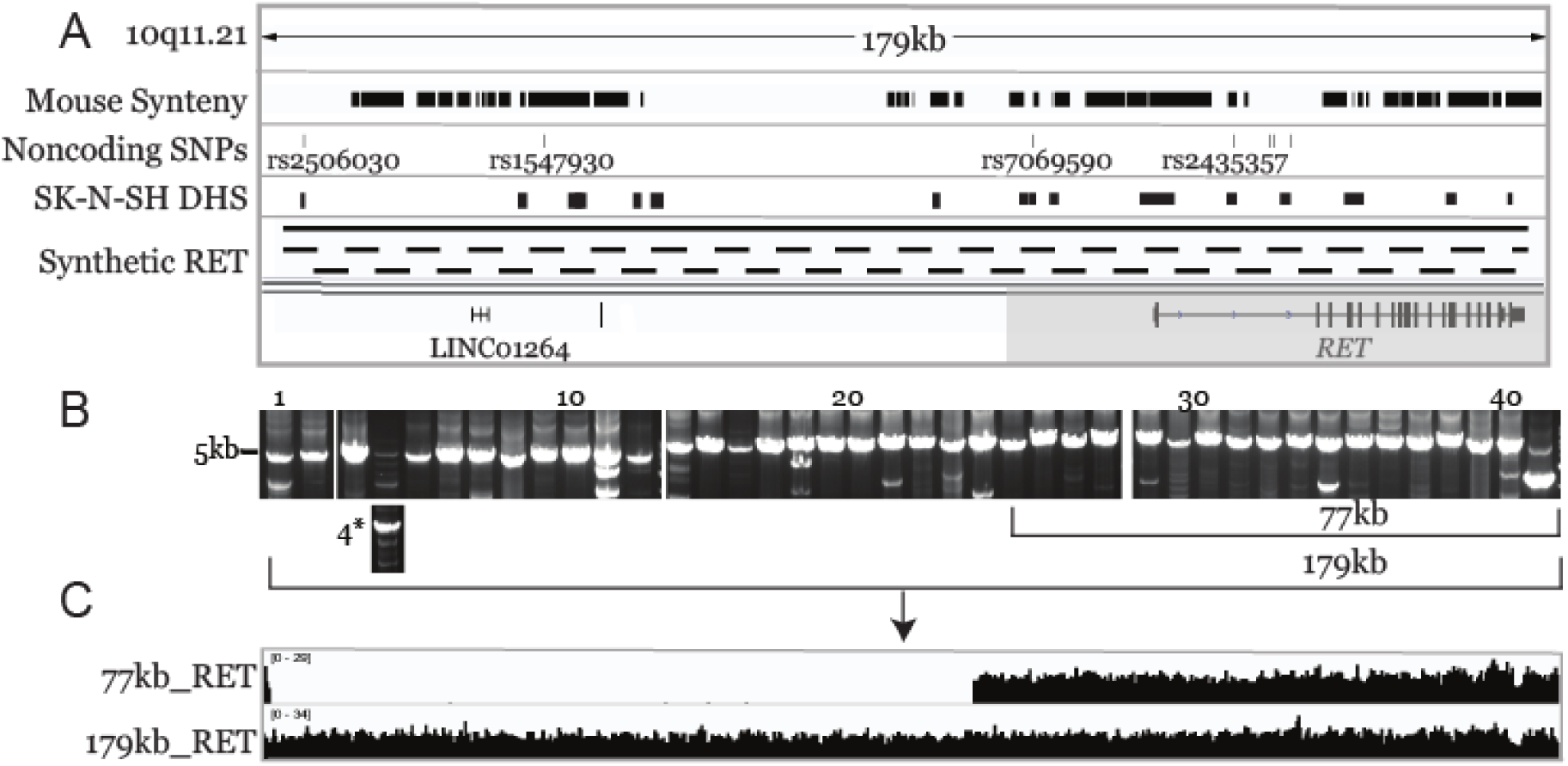
Synthetic design and construction of the human *RET* locus. **(A)** Genomic snapshot of the *RET* locus on human chromosome 10 (chr10:42951900-43130851, hg38) annotated with the Hirschsprung disease risk variants, regions of mouse synteny and DNaseI hypersensitive (DHS) sites in the neuroblastoma cell line SK-N-SH. **(B)** EtBr-stained agarose gel image of ∼5kb PCR fragments use to assemble the 77kb *RET* locus in yeast. **(C)** Whole genome sequence coverage of the primary 77kb *RET* locus and largest 179kb locus synthesized in yeast from PCR products in (B).

To construct this synthetic locus, we segmented the region into ∼5kb lengths with 250bp overlaps and designed PCR primers (Table S1) using the MenDEL software (**Methods**). The 5kb segments were initially generated by PCR from bacterial artificial chromosomes using a high-fidelity polymerase and an optimized PCR program (**Figure 1B)**. Segment 4 was in particular very GC rich and required addition of 1M Betaine to construct. Two additional 500bp fragments homologous to the *RET* locus and the plasmid backbone were synthesized as Gblocks from IDT (**Table S1**). After synthesis, the fragments were purified, pooled in equimolar amounts, and transformed into yeast using the standard LiAc transformation method along with a BsaI-linearized dual *S. cerevisae/E. coli* “YAV” (yeast assembly vector) backbone; our goal was to utilize the endogenous yeast homologous recombination pathway to generate an intact delivery vector. We then screened colonies for successful recombination using highly automated PCR with junction primers that spanned the 500bp overlapping regions between any two fragments in the assembly. Despite several attempts, no YAVs bearing the entire 179kb locus could be synthesized in this manner. However, a colony bearing the majority of the 77kb *RET* locus was obtained (**Table S4**). This required further editing with CRISPR/Cas9, and subsequent yeast recombination, to replace 3 missing segments of DNA with the final YAV confirmed to be the entire desired locus by whole genome sequencing (**Figure 1C)**. To construct the entire 179kb locus, we utilized gRNAs to digest fragments of ∼40kb and ∼60kb in length from 2 BACs *in vitro* and transformed these into yeast with a Cas9 linearized version of the 77kb delivery YAV. This successfully yielded an entire intact 179kb *RET* locus in yeast, which was subsequently recovered into *E. coli* (**Figure 1C**).

To deliver these large *RET* constructs into mice, we utilized a BxbI recombination system to directly insert the locus into mouse zygotes by co-injecting purified delivery vector along with BxbI integrase mRNA (**Figure 2A**).^13^ This system makes use of heterologous (GA) and (GT) attB/attP recognition sequences in the donor vector and the *Rosa26* locus, respectively, to deliver the DNA in a directional manner (**Figure 2B**). We spent considerable time adjusting buffer and delivery conditions from the primary laboratory site to the injection site to achieve successful integration via this method. Ultimately, the successful strategy was purification of the vector in TE buffer with addition of polyamines (spermine, spermidine and NaCl, see Methods), maintenance at 4°C on wet ice, followed by microinjection within 1 week of delivery between the two physical laboratory locations utilized in this work. From condition 1 (0.5ng/µl Donor), 38 mice were born from a total of 155 embryos transferred (25% survival) and no candidates were identified with the correct insertion. From condition 2 (30ng/µl Donor), 54 mice were born from a total of 156 embryos transferred (35% survival) and a single candidate was identified with the correct insertion (2%). Evidence of off-target integration (i.e., random integration) was present in mice from both conditions (5/38, 13% from Condition 1 and 12/54, 22% from Condition 2). We confirmed successful integration of the entire *RET-s* locus in a female germline transmitted N1 progeny by whole genome sequencing (**Figure 2C)**.

**Figure 2.**
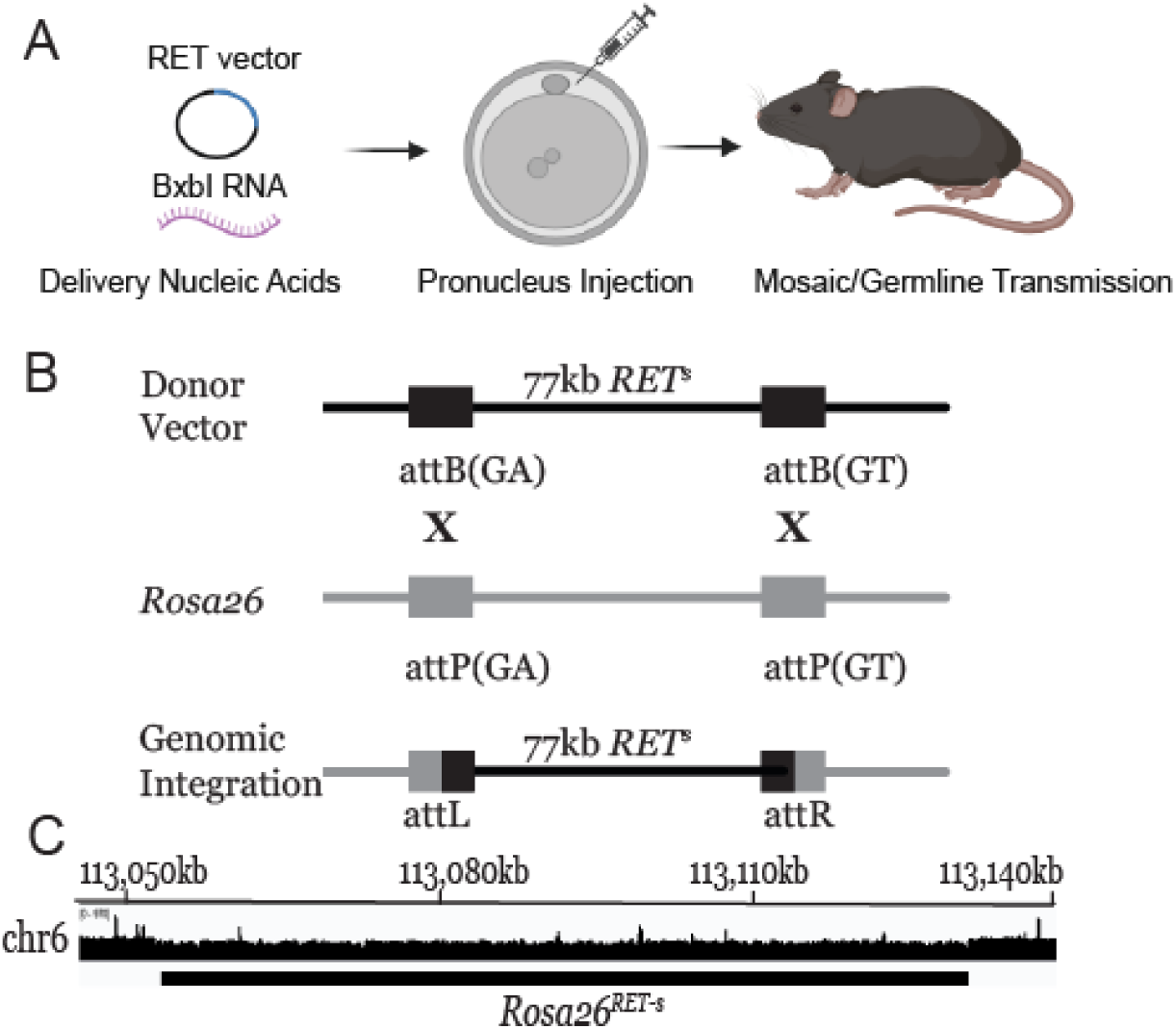
BxbI targeted DNA delivery into mouse zygotes. **(A)** Donor DNA and BxbI RNA are directly injected into mouse zygotes. **(B)** Donor DNA is integrated through use of two heterologous attP (48bp) and attB (38bp) sites by the bacteriophage serine integrase BxbI into the *Rosa26* locus leaving behind only 43bp genomic attL/R scars.^13^ **(C)** Whole genome sequence coverage of the initial germline transmitted mouse bearing the 77kb HSCR sensitive haplotype *RET-s*.

We took advantage of our initial mouse line bearing both the *RET-s* locus within *Rosa26* and the endogenous *Ret* locus to compare their relative transcription levels in a homozygous background (**Figure 3A**). To do so, we collected RNA from 6 tissues at birth (P0): heart, lung, stomach, small intestine, colon, and kidney. Heart and lung were considered negative controls with minimal to no *Ret* expression. We found that in nearly all cases, except for the heart, *RET-s* expression was ∼10% of mouse *Ret* levels in key developmental tissues of the gastrointestinal tract and kidneys and with no detectable difference between male and female mice (**Figures 3B and C**). To ensure that the expression we were observing was not artefactual, we tested our TaqMan probes for specificity to the human and mouse genes using human CHP212 and mouse Neuro2a neuroblastoma cell lines, respectively (**Figure S1**). We also performed PCR, utilizing primers across all of the 20 *RET* exon-intron junctions on cDNA synthesized from RNA isolated from mouse colon of a P0 sample. Sanger sequencing confirmed that all introns were correctly spliced from *RET* and all exons were present (**Figure S2**).

**Figure 3.**
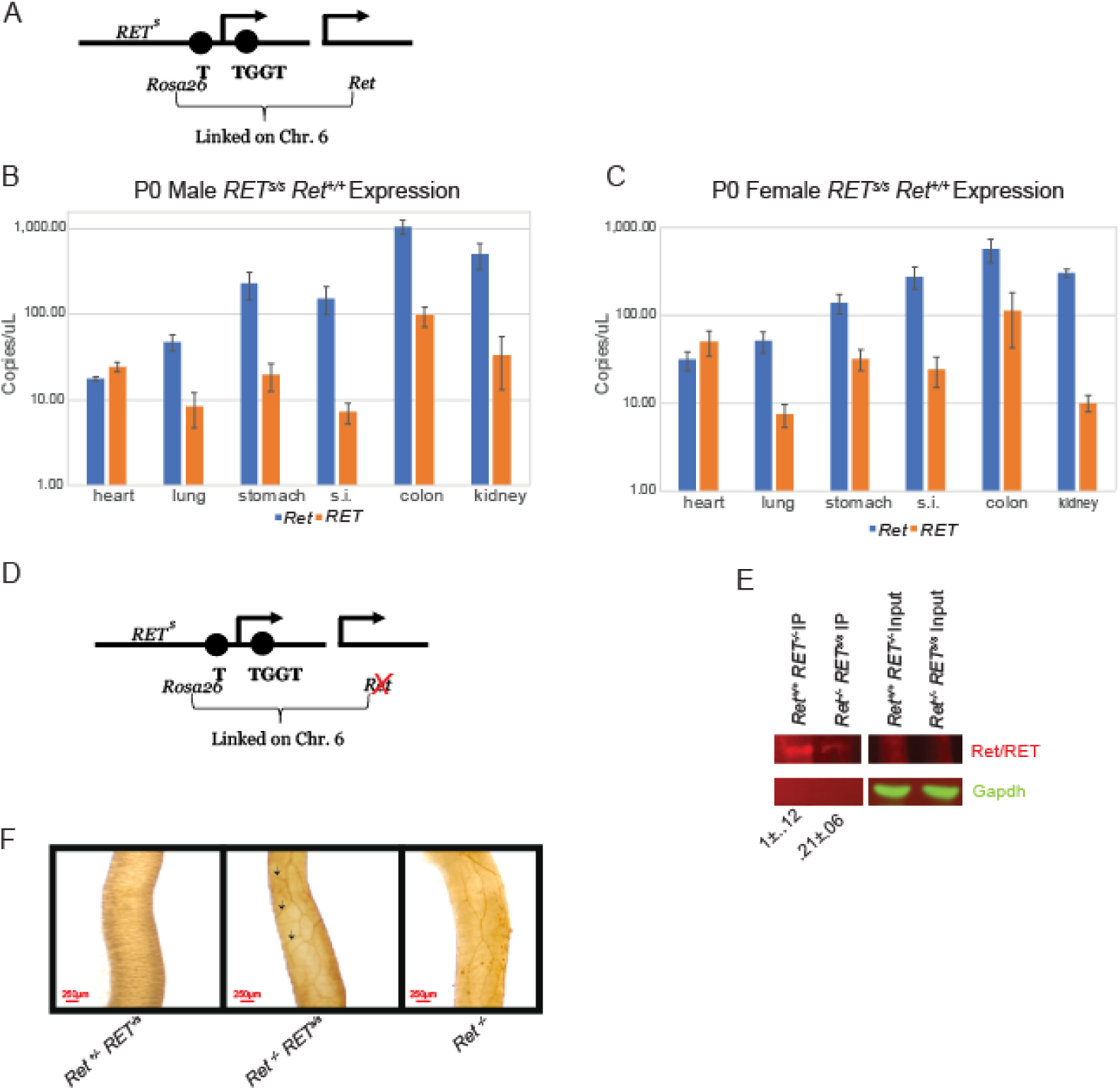
Human *RET* expression relative to mouse *Ret*. **(A)** Schematic of the mouse genotypes used in **(B, C)** in animals in which both mouse *Ret* and human *RET* are present. **(B, C)** ddRT-qPCR of *RET* and *Ret* expression within selected tissues in male and female mice. Error bars represent standard error of 4 biological replicates. **(D)** Schematic of mouse genotypes used in Figure (E) in which mice are solely dependent upon *RET* or *Ret*. **(E)** Representative western blot of RET and Ret expression from immuno-precipitated colon tissue of P0 mice. Expression is quantified relative to mouse *Gapdh* and quantified from 3 biological replicates. **(F)** Acetylcholinesterase staining of AChE positive neurons (dark brown) in the distal colon of WT (left), *RET-s* (middle), and *Ret* null mice (right). Scale bar (250 μM)

These results gave us confidence that the human *RET-s* gene was intact and was being correctly transcribed and spliced albeit at low levels. Consequently, to study the effects of the *RET-s* gene alone, we proceeded to delete exons 2-6 of the endogenous *Ret* gene using CRISPR-Cas9 directed editing with 2 gRNAs directly in zygotes. We performed this modification in both a mouse line bearing *RET-s,* which is in cis with *Ret* on chromosome 6, as well as an isogenic WT line. We confirmed that deleting exons 2-6 of *Ret* in the isolated line was sufficient to eliminate detectable levels of *Ret* RNA as well as protein expression (**Figure S3.**) We then performed immuno-precipitation followed by western blot detection of RET at P0 from *Ret^-/-^ RET^s/s^* colon and found that it was expressed 4.7-times lower than *Ret* in age matched *Ret^+/+^* samples (**Figure 3E**).

Given that the expression of *RET-s* was <50% of *Ret* at both the transcriptional and translational levels, we expected that *Ret*^-/-^ *RET*^s/s^ mice would exhibit classic hallmarks of HSCR based on our earlier studies.^13^ Notably, these *Ret*^-/-^ *RET*^s/s^ mice do not survive beyond P0 and succumb to kidney aplasia (data not shown). We examined these mice for presence of the ENS by acetylcholinesterase staining of AChE+ neurons. In compound heterozygous *Ret^+/-^ RET^-/s^* mice where *Ret* is present, the colon is innervated at WT levels as expected (**Figure 3F**). Upon introduction of a second copy of *RET-s* and depletion of *Ret,* we see extensive aganglionosis and elongated AChE+ neurons throughout the intestinal tract up to the stomach (**Figure 3F**). However, qualitatively there appears to be a slight increase in neuronal branching (black arrows) compared to the phenotype in the *Ret^-/-^* mice in which we deleted exons 2-6, suggesting that the 21% of expression does have some positive impact on neuronal innervation (**Figure 3F)**.

We hypothesized 2 possibilities for the low expression levels of *RET-s* expression. One, perhaps the *RET-s* locus is recognized as a foreign source of DNA and thus actively repressed through DNA methylation or heterochromatinization of *RET-s*. To examine this, we utilized Oxford Nanopore long read sequencing to both reconfirm presence of *RET-s* within *Rosa26* and examine methylation levels of the DNA by adaptive sampling of ∼1% of the genome in *Ret^+/+^ RET^s/s^* mice. We found that 536/745 CpG sites (71%) exhibited methylation within the *Ret* locus whereas *RET-s* exhibited 895/1556 methylated sites (57.5%) when requiring at least 50% of reads to call the site as methylated. This suggested to us that DNA methylation levels were normal within *RET-s*. We further examined *RET-s* chromatin with ATAC-seq and found that all of the open chromatin peaks present from 2 WT human colon samples in ENCODE^14^ were found in samples prepared from *Ret^-/-^ RET^s/s^* mouse colon (**Figure 4A**). Collectively, these results lead us to believe that the majority if not entirety of the *RET-s* locus is not being silenced.

**Figure 4.**
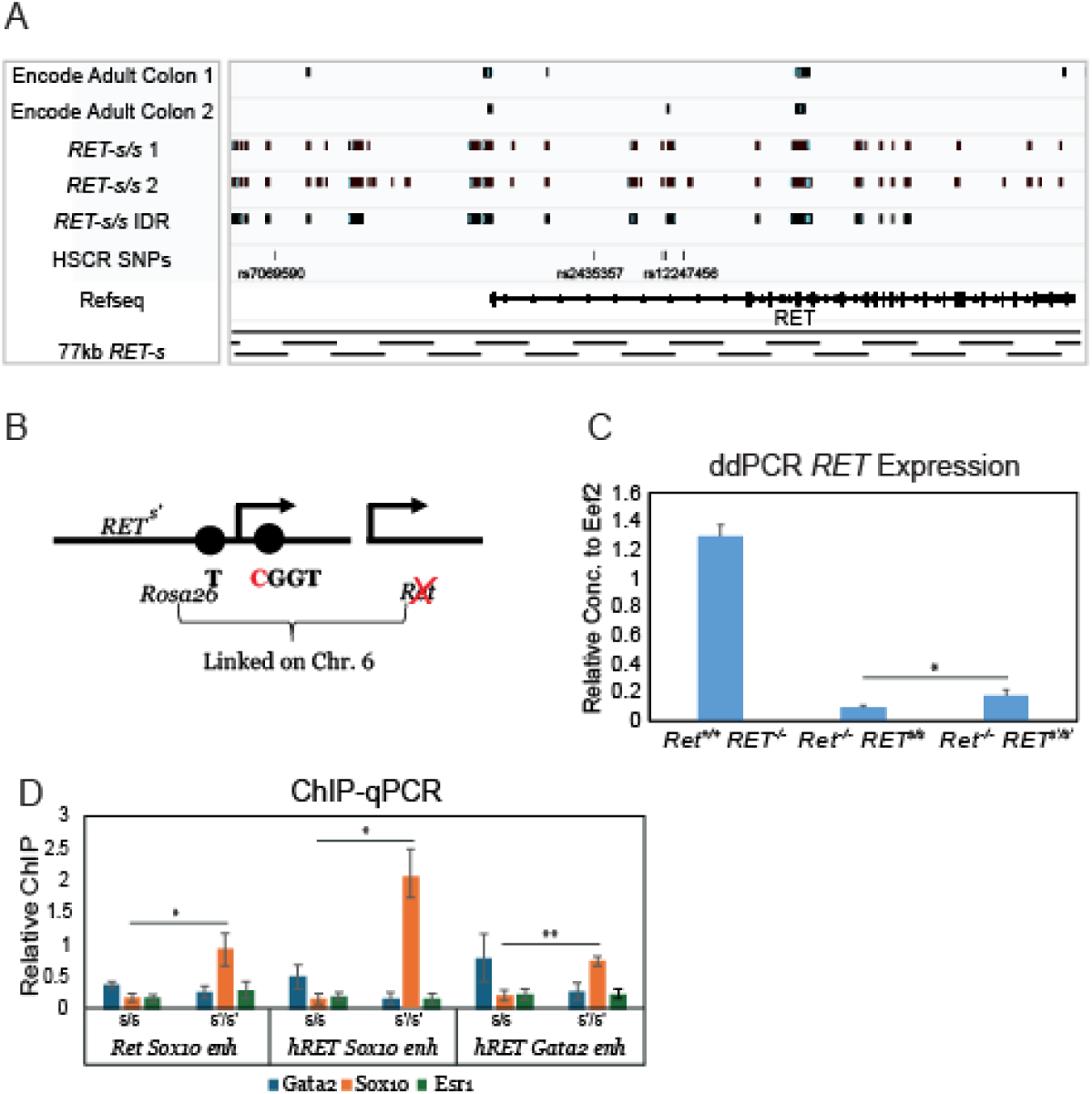
*RET-s* is not heterochromatized and its expression can be increased by introduction of a resistant SNP. **(A)** ATAC-seq narrowpeak tracks from 2 ENCODE human colon samples as well as 2 *RET-s* E14.5 mouse colon samples. The IDR (Irreproducible Discovery Rate) track represents peaks that were reproduced by the 2 biological replicate *RET-s* samples with a false positive cutoff of 5% **(B)** Schematic showing which SNP was edited for the *RET-s’* mice. **(C)** ddPCR of *Ret* vs *RET* expression in 3 different homozygous mouse lines from P0 colon tissue. *p<.05 two-tailed t-test **(D)** ChIP-qPCR of E14.5 colon tissue for 2 homozygous mouse lines for 3 different transcription factors. *Esr1a* was used as a control. Signal is relative to input control DNA before IP. *p<.05 **p<.01 two-tailed t-test.

Our second hypothesis for the lack of *RET-s* expression is that it is by definition a *sensitive* haplotype and we have simply crossed the threshold beyond which *Ret* can be rescued. To test this, we edited the Sox10 enhancer binding site, rs2435357, from the sensitive “T” to resistant “C” allele by CRISPR/Cas9 editing in mouse zygotes and call this line *RET-s’* (**Figure 4B**). We confirmed successful editing by Sanger sequencing of the allele from the initial germline transmitted line. We then examined expression of *RET-s’* relative to WT *Ret* and *RET-s* within the colon and find that there is a roughly 1.9-fold increase in *RET* expression at P0 (Figure 4C). This is similar to data from another mouse line in our lab in which deletion of the conserved Sox10 enhancer in *Ret* resulted in an approximately 5% reduction of *Ret.*^16^ Additionally, we find that editing the “T” to “C” allele resulted in a significant increase in Sox10 binding to the *RET-s’* Sox10 enhancer site assayed by chromatin immunoprecipitation from E14.5 intestinal tissue (Figure 4D). We also note an increase in Sox10 binding at both the conserved *Ret* Sox10 site as well as the Gata2 enhancer binding site located further upstream and thought to be in contact with the Sox10 enhancer in 3D space (Figure 4D). Collectively, we believe our results suggest that the *RET-s* locus is essentially a minimally active and intact human locus that exhibits diminished *RET* expression at least in part due to presence of 5 sensitive SNPs within critical enhancer elements. Future work will be required to increase the size of the locus as well as observe further increases in *RET* expression upon conversion of the remaining 4 SNPs to their protective states to determine their individual and combined effects.

## Discussion

Our studies demonstrate a novel approach to construct mouse models of complex human genetic disorders by inserting a large human gene encoding the disease associated haplotype of the *RET* locus directly into the mouse genome at a safe harbor locus, *Rosa26*. This model enables comprehensive studies of both the human coding gene and its cis regulatory elements, the latter being the site of the disorder’s sensitive variants. Although generating the relevant DNA constructs in budding yeast and *E. coli*, even as large as 179kb, is straightforward, insertion of a single, large, engineered segment of DNA directly into a specific target into mouse zygotes is both challenging and suffers from low efficiency. To date the most promising approach appears to be “mSWAP-IN”^17^ technology in which human loci greater than 100kb in length can be integrated into the genome by repeated delivery of genomic fragments with theoretically no limit to the number of deliveries. Furthermore, 200kb in one step with this approach may become common place with at least one locus showing early success (unpublished data from the JDB lab).

Our current efforts led to a humanized mouse model of HSCR by the direct insertion of a 77kb *RET* haplotype construct into the mouse genome using Bxb1 integrase. This is a significant technical advance since previously published work utilizing the same delivery procedure produced 30kb insertions.^13^ The technology used also bypasses many of the limitations of traditional genesis methods, allowing for the precise inclusion of critical noncoding regulatory elements, which are often the primary drivers of complex traits. The fact that the human *RET* locus was functionally expressed in the mouse, with all introns correctly spliced, validates the model as a powerful tool for studying human gene regulation in a living system.

However, the findings also highlight a critical limitation: despite the model’s technical success, the low expression level of the human *RET* gene (21% of mouse *Ret* levels) and the resulting full HSCR penetrance in homozygous animals suggest that our understanding of *RET* gene regulation is still incomplete. This discrepancy implies the necessity of the regulatory elements located 100kb further upstream of the current locus. It is also possible that trans-acting factors in the mouse fail to appropriately interact with the human *RET*, including TFs, epigenetic regulators, or even lack of presence of RET’s coreceptor, GFRA1, and ligand, GDNF. While we considered that the human gene may simply express lower than the mouse gene, the TPM values of *RET* (1.1-1.6) from public sigmoid and transverse colon data in the human genome atlas and *Ret* (1.6) from our P0 colon datasets suggests this isn’t the case. This is a significant finding because it challenges the assumption that simply transplanting a human gene and its known regulatory regions into a mouse is sufficient to recapitulate human gene expression. Thus, the 179kb locus, which we successfully constructed but have not yet inserted, would be the next logical step. It contains more of the implicated SNPs and enhancers, and its insertion may yield a different expression pattern. Given the lack of current understanding, perhaps ideal candidate genes are those that are shorter in length, which are generally highly expressed such as housekeeping genes^18^, and often regulate longer genes that may be associated with specific and essential biological processes such as embryonic development^19^ or neuronal processes^20^.

The extensive loss of enteric nervous system (ENS) ganglia in our *Ret^-/-^ RET^s/s^* mice, despite low-level expression of the human *RET* gene, supports our recent data on HSCR pathogenesis. Earlier mouse studies focused on high-penetrance, loss-of-function mutations, but this model provides a tangible link between a hypomorphic allele and a severe disease phenotype. The human *RET-s* allele, even at a mere 21% of WT levels, is not sufficient to rescue the *Ret* knockout phenotype. This suggests that a critical threshold of RET protein expression is necessary for normal ENS development, and falling below this threshold, even slightly, leads to complete aganglionosis. We recently suggested this threshold begins around 30% of the combined loss of *RET* and *EDNRB* gene expression due to their epistatic nature.^14^

This finding also offers a mechanistic explanation for the highly variable penetrance seen in human HSCR patients. The phenotype in our model suggests that the accumulation of low-risk hypomorphic alleles in a single gene, or a combination of them, can collectively push an individual’s *RET* expression below the necessary developmental threshold, leading to the disease. The slight increase in neuronal density observed qualitatively in the *Ret^-/-^ RET^s/s^* mice compared to the *Ret^-/-^* mice, while not sufficient to prevent the disease, hints at a subtle dosage effect, further supporting the idea of a critical threshold. This mouse model is therefore an ideal tool to systematically test how different combinations of human SNPs in the *RET* locus contribute to the ultimate expression level and disease phenotype and whether or not we can rescue this phenotype by creating haplotypes with increasing numbers of resistant enhancer alleles.

The successful creation of this humanized mouse model opens up a new avenue for studying the functional consequences of complex genetic variation. Namely, therapeutic intervention for a genetic disorder in which we know the root cause. *RET* variants are found in nearly 50% of HSCR cases, suggesting a large target population in need of an alternative to surgery, which although lifesaving is not without its complications. Aside from increasing the size and complexity of the model, we envision that this model can serve as a platform to trial *in utero* gene therapies aimed at activating *RET* either at the transcriptional or translational levels. Ultimately, this work is not just about understanding HSCR; it is a foundational step towards mouse models that can accurately reflect the effects of human genetic variation and treatment of complex diseases.

## Materials and Methods

### Mouse Strains

We used the C57Bl/6J strain for all of the work described here and all strains are available from the Jackson Laboratory. The humanized 77kb *RET^s^* allele inserted into *Rosa26* in a *Ret^+/+^* background (C57BL/6J-Gt(ROSA)26Sor^em1^(RET)^Arz^/Arz) is denoted strain JR37659. The isolated *Ret* null allele (C57BL/6J-Ret^em1Arz^/Arz) and the cis *Ret* null allele (C57BL/6J-*Gt(ROSA)26Sor^em^*^1^(RET)*^Arz^ Ret^em2Arz^*/Arz) in the humanized *RET* genetic background were made by targeting Cas9 to *Ret* exons 2-6 with a dual guide approach and are denoted as strains JR38783 and JR38676, respectively. Finally, the strain with the Sox10 enhancer disease variant site mutated to the resistant allele (C57BL/6J-Gt(ROSA)26Sor^em30^(RET*)^Arz^ Ret^em2Arz^/Arz) is denoted as JR40104. The gRNA sequences used for creating this mutation are provided in Table S1. All procedures were approved by the New York University IACUC (protocol number: IA17-01779).

### Mouse Genotyping

DNA was extracted from ear punch biopsies or tail snips (postnatal day 0 mice) using a Proteinase K digestion buffer (50mM Tris pH 7.5, 5mM EDTA, 1% SDS, 0.2M NaCl, 0.2mg/mL Proteinase K) for 3 hours at 55°C followed by 1:1 phenol: chloroform extraction and ethanol precipitation. All PCRs were done in a single reaction using Go-Taq Green master mix with a universal primer and two allele specific primers. Primers are listed in Table S1. The WT *Rosa26* PCR product is 700bp while the human *RET* gene is 400bp. The PCR product for the WT *Ret* allele is 120bp. For the isolated *Ret* null allele, the PCR product is 250bp and 430bp for the *Ret* null allele that is in *cis* with the human *RET* gene on chromosome 6.

### Synthetic *RET* design

The 179kb variant of the synthetic *RET* locus was designed to include all noncoding DNA up to 500bp beyond a previously defined RARB enhancer binding site^6^ and the entire *RET* coding sequence plus 500bp beyond the 3’UTR, as annotated in Refseq; this corresponds to hg38 chr10:42951900-43130851. The 77kb *RET* locus primarily used in this work to generate mice includes all noncoding DNA just beyond a previously defined GATA2 enhancer site^6^ as well as the coding sequence plus 500 bases beyond the 3’UTR annotated in Refseq; this corresponds to chr10:43053532-43130851. We used MenDEL software^21^ to split the synthetic locus into smaller DNA segments of approximately 5kb each for PCR synthesis with 500bp of overlap to enable yeast assembly (described below). PCR primers for both construction and junction overlap were designed by MenDEL and are available in Table S1.

### Synthetic yeast *RET* assembly and sequence validation

All primers together with yeast and *E*. coli strains utilized in construction are provided in Tables S1, S2, and S3, respectively. The primary 77kb human *RET* locus was initially constructed by yeast transformation of the By4741 strain^22^ grown to log phase in YPD media using the lithium acetate transformation method.^23^ A BsaI linearized vector backbone, pLM1050, along with 18 PCR fragments derived from the bacterial artificial chromosome RP11-419K10^24^ were transformed into yeast and grown on synthetic complete media missing leucine. pLM1050 allows for both yeast auxotrophic selection via the *LEU2* gene and bacterial kanamycin antibiotic selection via the *KanR* gene. Initial transformants were screened with primers spanning the approximately 500bp junction overlaps between PCR segments. Crude yeast gDNA was prepared by boiling in 20mM NaOH at 98°C for 3 min, followed by cooling at 4°C for 1 min three times. A Labcyte Echo 650 liquid handler was used to dispense 20 nL crude gDNA and 10 nL premixed junction primer pairs (50 µM) into a LightCycler 1536 Multiwell Plate (Roche) containing 1 µl 1× LightCycler 1536 DNA Green mix (Roche). Successful yeast artificial vectos (YAVs) were identified based on Ct values less than 30 for all junctions. A water control was included to filter out nonspecific junction pairs. Candidate assemblies were verified by whole genome sequencing on a Miseq sequencer. Libraries were prepared from 100 ng of DNA isolated with a Zymogen YeastStar Genomic DNA kit using the NEBNext Ultra II FS DNA Library kit, according to manufacturer’s instructions for each.

The 179kb hRET locus was constructed in yeast by using Cas9 (NEB) to digest a ∼60kb fragment of DNA from 1 µg of RP11-124O11 BAC and 40kb from 1 µg of RP11-419K10 for 4 hr at 37°C. Cas9 was also used to induce a double strand break in the upstream region of the 77kb hRET YAV pRDF90 using the same conditions. After digestion, 500ng of each digest was combined with 5 µL of PCR product of junction overlapping IDT-synthesized gene blocks in a standard LiAc yeast transformation. Colonies were selected on SC media lacking leucine for two days at 30°C. Colonies were screen for the insert by colony PCR. Candidate clones were sequenced as previously described. The 179kb version was never utilized for mouse zygote injection due to limited initial success with the smaller 77kb variant, but is available for future attempts (**Table S3**.)

### Yeast Lithium Acetate transformation

An O/N yeast culture was used to inoculate a fresh 20 mL culture to a final OD₆₀₀ of 0.15, typically 200 μL. The culture was incubated at 30°C with rotation for two cell doublings, 3-4 hours, in YPD. Following incubation, cells were pelleted at 3000 rpm for 3 minutes and the supernatant was discarded. The cells were washed twice: first with 20 mL of sterile water and then with 20 mL of sterile 0.1 M lithium acetate (LiOAc). The final pellet was resuspended in 200 μL of 0.1 M LiOAc which is sufficient for 4 transformations. To perform the transformation, a master mix containing 29% polyethylene glycol (PEG), 0.1 M LiOAc, 0.7 μg/μL boiled salmon sperm, the yeast, and 10-20μL of DNA was added to a total reaction volume of 360 μL. The transformation was mixed by minimal pipetting to reduce shearing of large DNA constructs.

The transformation mixture was incubated at 30°C for 30 minutes then subjected to heat shock at 42°C for 15 minutes. Following the heat shock, the cells were pelleted by centrifugation at 3000 rpm for 3 minutes at room temperature and the supernatant was removed. The pellet was washed in 5 mM CaCl₂ then resuspended in 400 μL of 5 mM CaCl₂, mixed by pipetting, and incubated for 10 minutes at room temperature. The transformed cells were plated onto the appropriate selective medium. To ensure a sufficient number of colonies, we plated 50 μL and 350 μL volumes on separate plates. The plates were incubated at 30°C for 2 days and utilized for colony PCR, purification, and eventual DNA isolation.

### Recovery into *E. coli* and Purification

A Zymoprep Yeast Plasmid Miniprep II kit was used to recover YAV DNA and electroporated into TransforMax EPI300 Electrocompetent *E. coli*. After a 1hr recovery at 30°C, bacteria were plated on *Kanamycin* selection plates for overnight growth at 30°C. Colonies were screened by colony PCR of the edge junctions for retention of the insert and/or BamHI restriction enzyme digestion of a 5 mL alkaline lysis miniprep. Successful colonies were grown in 5 mL starter cultures overnight then grown in 300 mL primary cultures induced with 0.04% L-Arabinose for 4 hours. DNA was isolated using an Nucleobond Xtra BAC DNA isolation kit from TakaraBio according to manufacturer’s instructions except that the: 1) DNA was chilled at 4°C after addition of resuspension buffer for 20 minutes to induce supercoiling, 2) Cellular debris was pelleted after addition of neutralization buffer and the supernatant was directly applied to the column, and 3) DNA was precipitated in low-bind DNA Eppendorf tubes and pooled in 100 μL of TE buffer. Generally, yields that were greater than 100 ng/μL resulted in success for downstream engineering applications. Fresh RNAseA was added to the lysis mix for preps used in zygote injections.

### Mouse Zygote Injection

The *RET-s* mouse was generated at The Jackson Laboratory using Bxb1 Integrase Mediated Cassette Exchange (B-RMCE) as previously described.^13^ Using heterozygous zygotes of the *Rosa26* Dual Bxb1 Landing Pad mouse strain (JAX Stock #36152), B-RMCE of the gene was achieved following pronuclear microinjection of commercially prepared (TriLink BioTechnologies) Bxb1 Integrase mRNA (100 ng/µl), Donor Plasmid pRDF90-1 (two concentrations tested: 0.5 and 30ng/µl), and RNasin® Ribonuclease Inhibitor (Promega, cat# N2515; at 0.2U/µl). pRDF90-1 was shipped at 4°C on wet ice in BAC buffer supplemented with polyamines (10 mM Tris-HCl, pH 7.5, 0.1 mM EDTA, 30 µM spermine, 70 µM spermidine, 100 mM NaCl). Both Bxb1 mRNA and the Donor Plasmid were ultra-purified for microinjection using phenol-chloroform extraction and ethanol precipitation. All reagents for microinjection were prepared in nuclease-free BAC Buffer supplemented with polyamines (10 mM Tris-HCl, pH 7.5, 0.1 mM EDTA, 30 µM spermine, 70 µM spermidine, 100 mM NaCl). Microinjected zygotes were transferred to pseudo-pregnant females and carried to term. A single candidate (Male, #8763) identified was bred to wild-type C57BL6/J females (JAX Stock #000664) to verify germline transmission (GLT) and establish N1 generation heterozygotes. After these offspring were confirmed to be free from off-target integration events, they were used to establish the colony, now designated with the JAX Stock Number: #037659 C57BL/6J-Gt(ROSA)26Sor<em1(RET)Arz>/Arz (*RET-s*).

### Mouse Whole Genome Sequencing

DNA was extracted from ear punch biopsies as previously described^14^ and libraries were prepared using an Illumina Truseq DNA PCR-free kit with an average insert size of 350 bp. Libraries were sequenced on an Illumina Hiseq instrument. Reads were filtered using standard metrics and then mapped to a custom mouse reference genome bearing the *RET* sequence inserted within the *Rosa26* locus. Raw reads of the initial founder mouse for *RET-s* are available upon request.

### Oxford Nanopore Technologies Sequencing

High-molecular-weight (HMW) DNA was isolated from tissue using the Monarch HMW DNA Extraction Kit (New England Biolabs) following the manufacturer’s protocol with an agitation speed of 2000 rpm. DNA quantity and quality were assessed using a NanoDrop 2000 spectrophotometer (Thermo Scientific), Qubit 3.0 dsDNA BR Assay (Thermo Scientific), and Femto 165 kb Assay (Agilent Technologies). Libraries were constructed using the Ligation Sequencing Kit v14 (Oxford Nanopore Technologies) with third-party reagents. Briefly, 5’ DNA ends were dephosphorylated before the addition of Cas9 ribonucleoprotein particles (RNPs) containing bound crRNA and tracrRNA to cleave the region of interest (ROI). Cas9 cleavage generated blunt ends with ligatable 5’ phosphates. Following dA-tailing of all DNA, sequencing adapters were ligated primarily to the Cas9 cleavage sites. Library quality and concentration were verified using the Genomic DNA ScreenTape (Agilent Technologies) and Qubit dsDNA BR Assay (Thermo Fisher Scientific). Sequencing was performed on an Oxford Nanopore GridION device. Basecalling was conducted using Dorado v7.3.11.

### ATAC-sequencing

Nuclei were isolated from the intestinal tract of 4 fresh E14.5 embryos using a 10x Genomics Nuclei Isolation kit PN-1000493 according to manufacturer’s instructions and pooled into two samples. ATAC-seq libraries were prepared with an Active Motif kit (53150) using 100,000 nuclei as input. Libraries were sequenced on an Illumina Novaseq to a depth of 50M paired end reads. Reads were filtered for poor quality, duplication, and mapped to hg38 by utilizing the ENCODE ATAC-seq pipeline available online (https://github.com/ENCODE-DCC/atac-seq-pipeline). ENCODE samples ENCFF217BRO and ENCFF396QHI were directly downloaded from the ENCODE online repository as filtered bigBED files.^14^

### Sanger Sequencing of *RET* cDNA

PCR was performed on cDNA from *RET^+/s^ Ret^+/+^* P0 colon samples used in ddPCR experiments with primers designed to span all exon-intron junctions of the human *RET* allele. PCR products were then cleaned with a Zymogen DNA Clean and Concentrator-25 kit. Eluted DNA were Sanger sequenced using Azenta Life Sciences services. All raw call sequences are available upon request.

### Acetylcholinesterase Staining

AChE staining was performed as previously described.^18^ In short, P0 mice were euthanized by CO2 exposure followed by decapitation. The intestinal tract was dissected and rinsed in cold phosphate buffered saline (PBS). Fecal matter was removed with mechanical force using tweezers and the tissue was fixed in a 4% formaldehyde PBS solution. Tissue was incubated in a 1.72M sodium sulfate solution overnight at 4°C. The tissue was then immersed in substrate buffer (0.2 mM Ethopropazine HCl, 4 mM Acetylthiolcholine iodide, 10 mM Glycine, 2 mM Cupric sulfate pentahydrate, 65 mM Sodium acetate) at room temperature. A 1.25% sodium sulfide pH 6.0 solution was then added to develop the stain for 5 minutes and rinsed in distilled water. Images were taken using a stereomicroscope.

### RNA Extraction and ddPCR

Gut tissue was dissected and rinsed in cold 1X PBS and immediately snap-frozen in liquid nitrogen. RNA was extracted with a Qiagen RNeasy plus kit according to the manufacturer’s instructions. RNA purity was assessed with a nanodrop. cDNA was made using a superscript IV synthesis kit (Thermo Fisher) with 1 µg of RNA as input. cDNA was mixed with primers and ddPCR supermix for probes without dUTP (Biorad). Oil droplets were created with a Biorad QX200 droplet generator and quantified with a QX200 droplet reader. Endogenous *Ret* mouse transcripts were directly compared to the humanized allele using TaqMan probes designed to be specific for each species-specific allele.

### Western Blot

Intestinal tissue from P0 mice were dissected and immediately frozen in liquid N2. Tissue was lysed with cold lysis buffer (10 mM Tris-HCL pH 8.0, 250 mM NaCl, 1 mMEDTA, 1% Triton X-100) with addition of 1X protease inhibitors (Sigma) and ∼250 µL volume acid washed glass beads for 30 seconds at 30Hz in a Qiagen TissueLyser II machine. Protein was quantitated with a Pierce Protein Assay Kit. After protein normalization between samples, volumes were brought up to 600 µL and a 1/10 volume fraction was kept as input. The remaining volume was combined with 8 µL of anti-RET antibody (C31B4, Cell Signalling) and allowed to incubate with rotation overnight at 4°C. 40uL of lysis buffer-washed Protein A agarose beads (Thermo Scientific) were then added to the solution and incubated for an additional 4 hrs. Cold lysis buffer was used to wash beads three times. Protein was eluted in SDS sample buffer with reducing agent by boiling at 95°C for 2 minutes then immediately placed on ice. Samples were loaded into a pre-cast Genscript ExpressPlus 4-20% PAGE gel and run at 120V for 45 minutes. Gels were transferred using a wet transfer apparatus at 80V for 2 hours at 4°C to a nitrocellulose membrane. The membrane was blocked with a 5% powdered milk solution for 30 minutes. After rinsing in PBST, primary antibodies for either Ret/RET (421R25, Thermo Scientific) or Gapdh (D16H11, Cell Signalling) were incubated overnight at 4°C. The blot was rinsed in PBST at least 3 times for 10 minutes followed by incubation with infrared secondary antibody (Licor) for 1hr at room temperature. Blots were again rinsed at least 3 times for 10 minutes and images were taken on an Amersham Image Quant 800 western imager. Desitometry was performed using ImageJ.

### ChIP-qPCR

We performed chromatin immunoprecipitation by first collecting E14.5 intestinal samples of either *Ret^-/-^ RET^s/s^* or *Ret^-/-^ RET^s’/s’^* mouse intestine, followed by extensive washing in cold 1x PBS, and subsequent crosslinking in a 1% formaldehyde solution in PBS for 10 minutes. Crosslinking was stopped by addition of 125mM glycine and incubated for 5 minutes. Samples were transferred to microfuge tubes and snap-frozen by addition of liquid N2. After genotyping 2 E14.5 samples were combined per biological replicate with each TF assayed in biological triplicate. Tissue was first rinsed and then lysed with addition of cell lysis buffer (10mM Tris, 1mM EDTA, 100mM NaCl, .2%Np-40, .5% SDS, 1X protease inhibitors). Samples were sonicated using a Q55 sonicator set at 25% amplitude with 4 rounds of 30s “ON” exposure and placed on ice for several minutes before subsequent rounds. Cell debris were then precipitated by centrifugation at 10k rpm in a microcentrifuge. A 1:20 aliquot of solution was taken as the input DNA fraction and frozen at -80°C. The remaining chromatin solution was incubated with 5uL of the following antibodies for the respective TFs Sox10 (Santa Cruz:sc374170), Gata2 (Abcam:ab109241), and Esr1 alpha (Epicypher:13-2012) and incubated O/N with rotation at 4°C. The chromatin solution was then incubated with 50uL of anti-protein G agarose coupled beads that were pre-washed in lysis buffer for 4 hours at 4C. Beads were washed with lysis buffer (10mM Tris, 1mM EDTA, 100mM NaCl, .2%Np-40), high-salt lysis buffer (500mM NaCl), and LiCl wash buffer (10mM Tris, 1mM EDTA, 250mM LiCl, .2% Np-40). After washing, an addition of 100uL of 1x TE buffer was added to both the IP samples and input samples and de-crosslinked O/N at 65°C. Samples were then gently spun down and DNA was purified using a Zymogen DNA Clean and Concentrator PCR kit. DNA was eluted in 50uL of Zymogen elution buffer. qPCR was performed in technical duplicate and biological triplicate utilizing an independent chromatin input sample to generate standard curves. Primers are available in Table S1.

## Acknowledgements

We thank the members of the Boeke lab for guidance on the synthetic construction of the human *RET* locus in different delivery formats beyond those described here. Special thanks are owed to the NYU Genome Technology Center (GTC) for genome sequencing on various Illumina platforms. The GTC is partially funded by support grant P30CA016087 from the NYU Cancer Center as well an NIH Center for Excellence in Genomic Sciences (CEGS) grant to J.D.B. (RM1HG009491). We also gratefully acknowledge the contribution by the Genome Technologies Service at The Jackson Laboratory for expert assistance with Oxford Nanopore Technologies sequencing. This project was primarily funded by startup funds awarded to Aravinda Chakravarti, NIH grant HD028088 to A.C., and R01CA265978 to Vishnu Hosur of The Jackson Laboratory. Jef Boeke is a Founder and Director of CDI Labs, Inc., a Founder of and consultant to Opentrons LabWorks/Neochromosome, Inc, a Founder of JATech, LLC, and serves or served on the Scientific Advisory Board of the following: CZ Biohub New York, LLC; Logomix, Inc.; Rome Therapeutics, Inc.; SeaHub, Seattle, WA; Tessera Therapeutics, Inc.; and the Wyss Institute. The remaining authors declare no conflicts of interest.

## References

1 Altenburg E, Muller HJ. The Genetic Basis of Truncate Wing,-an Inconstant and Modifiable Character in Drosophila. Genetics. 1920 Jan;5(1):1–59.

2 Visscher PM, Wray NR, Zhang Q, Sklar P, McCarthy MI, Brown MA, Yang J. 10 Years of GWAS Discovery: Biology, Function, and Translation. Am J Hum Genet. 2017 Jul 6;101(1):5–22.

3 Fisher, R. A. (1918). The correlation between relatives on the supposition of Mendelian inheritance. Transactions of the Royal Society of Edinburgh, 52(2), 399–433.

4 Khera AV, Chaffin M, Aragam KG, et al. Genome-wide polygenic scores for common diseases identify individuals with risk equivalent to monogenic mutations. Nat Genet. 2018;50(9):1219–1224

5 Badner, J. A., Sieber, W. K., Garver, K. L. & Chakravarti, A. A genetic study of Hirschsprung disease. Am J Hum Genet 46, 568–580 (1990).

6 Chakravarti, A., McCallion, A. S. & Lyonnet, S. in The Online Metabolic and Molecular Bases of Inherited Disease (eds David L. Valle et al.) (McGraw-Hill Education, 2019).

7 Tilghman, J. M. et al. Molecular Genetic Anatomy and Risk Profile of Hirschsprung’s Disease. N Engl J Med 380, 1421–1432 (2019).

8 Emison AJHG

9 Amiel, J. et al. Hirschsprung disease, associated syndromes and genetics: a review. J Med Genet 45, 1–14 (2008).

10 Chatterjee, S., et al. Enhancer Variants Synergistically Drive Dysfunction of a Gene Regulatory Network In Hirschsprung Disease. Cell 167, 355–368 e310 (2016).

11 Chatterjee, S. & Chakravarti, A. A gene regulatory network explains RET-EDNRB epistasis in Hirschsprung disease. Hum Mol Genet 28, 3137–3147 (2019).

12 Chatterjee, S., Karasaki, K. M., Fries, L. E., Kapoor, A. & Chakravarti, A. A multi-enhancer RET regulatory code is disrupted in Hirschsprung disease. Genome Res 31, 2199–2208 (2021).

13 Low, B. E., Hosur, V., Lesbirel, S. & Wiles, M. V. Efficient targeted genesis of large donor DNA into multiple mouse genetic backgrounds using bacteriophage Bxb1 integrase. Sci Rep 12, 5424 (2022).

14 Fine, R. D., Chubaryov, R., Fu, M., Grullon, G. & Chakravarti, A. Joint disruption of Ret and Ednrb transcription shifts cell fate trajectories in the enteric nervous system in Hirschsprung disease. Proc Natl Acad Sci U S A 122, e2507062122 (2025).

15 Consortium, E. P. An integrated encyclopedia of DNA elements in the human genome. Nature 489, 57–74 (2012).

16 Fries, L. E., Grullon, G., Berk-Rauch, H. E., Chakravarti, A. & Chatterjee, S. Synergistic effects of *Ret* coding and enhancer loss-of-function alleles cause progressive loss of inhibitory motor neurons in the enteric nervous system. *bioRxiv*. 10.1101/2025.01.23.634550 (2025)

17 Zhang, W., Golynker, I., Brosh, R. et al. Mouse genome rewriting and tailoring of three important disease loci. Nature 623, 423–431 (2023).

18 Urrutia A. O., Hurst L. D. The signature of selection mediated by expression on human genes. Genome Res. 13, 2260–2264. (2003).

19 Lopes I, Altab G, Raina P, de Magalhães JP. Gene Size Matters: An Analysis of Gene Length in the Human Genome. Front Genet. 12:559998 (2021).

20 Sahakyan A. B., Balasubramanian S. Long genes and genes with multiple splice variants are enriched in pathways linked to cancer and other multigenic diseases. BMC Genomics 17:225 (2016).

21 German, S., Mitchell, L. A., Vela Gartner, A., Fenyö, D. & Boeke, J. D. MenDEL: PCR Primer Design as Constrained Optimization Process. *bioRxiv*. 10.1101/2022.06.26.496474 (2022).

22 Brachmann, C. B. et al. Designer deletion strains derived from Saccharomyces cerevisiae S288C: a useful set of strains and plasmids for PCR-mediated gene disruption and other applications. Yeast 14, 115–132 (1998).

23 Gietz, R. D. & Schiestl, R. H. High-efficiency yeast transformation using the LiAc/SS carrier DNA/PEG method. Nat Protoc 2, 31–34 (2007).

24 Osoegawa, K., de Jong, P. J., Frengen, E. & Ioannou, P. A. Construction of bacterial artificial chromosome (BAC/PAC) libraries. Curr Protoc Hum Genet Chapter 5, Unit 5 15 (2001).

