## Supplemental Figures for "BxbI-mediated insertion of a 77kb human RET sensitive haplotype into the mouse genome to generate a humanized model of Hirschsprung disease"

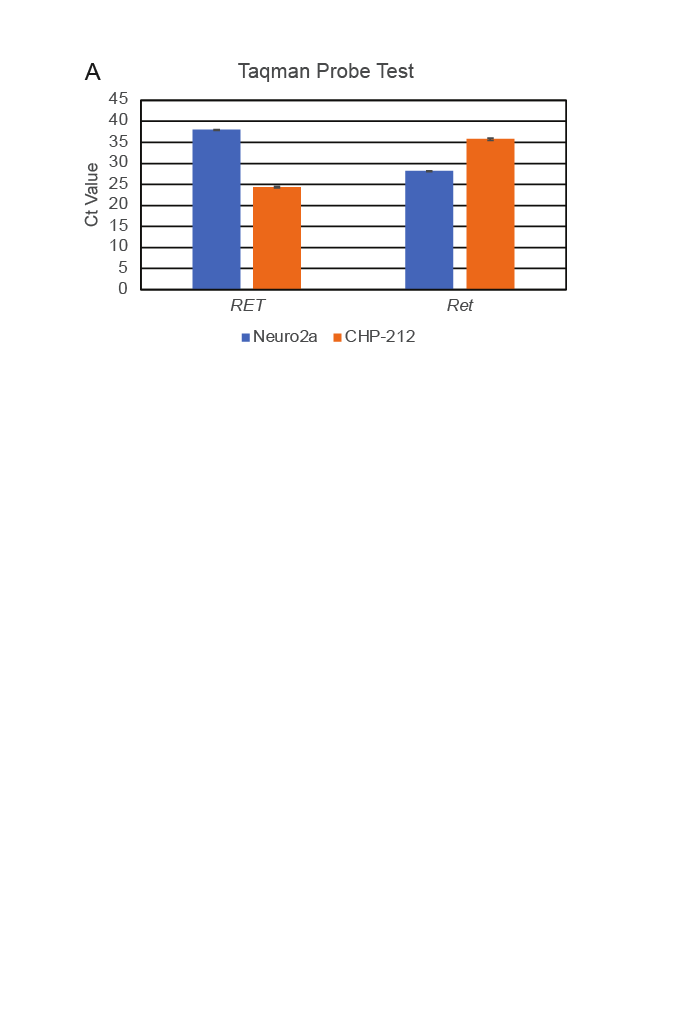

**Figure S1. Specificity of Taqman probes (A)** RT-qPCR experiment showing that the Taqman probes used for detecting mouse *Ret* and human *RET* gene expression are specific to the target organism as assessed in the neuroblastoma CHP212 (human) and Neuro2a (mouse) cell lines as proxies for specificity.

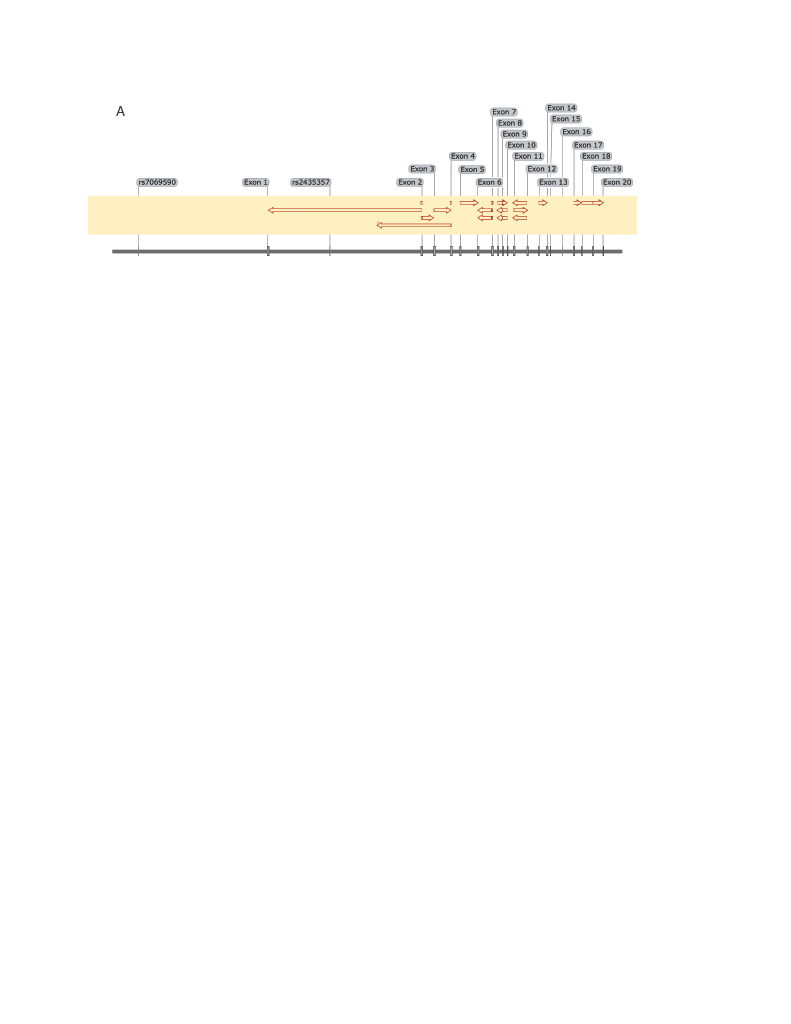

**Figure S2. Sanger Sequencing of *RET* cDNA (A)** SnapGene graphic showing Sanger sequencing alignment results from PCR products covering all annotated *RET* exon junctions.

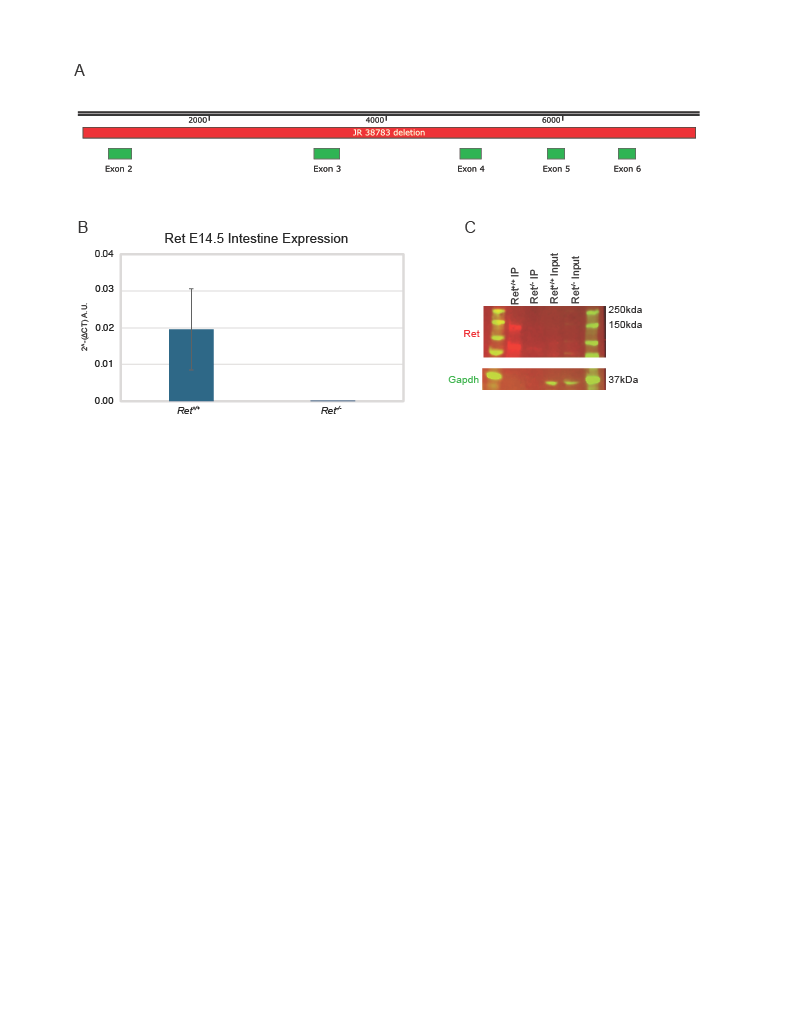

**Figure S3. *Ret* exon 2-6 deletion in the mouse (A)** SnapGene graphic showing the region of the *Ret* locus deleted in the isolated *Ret* deletion mutant. **(B)** RT-qPCR confirmation of loss of *Ret* expression in *Ret* exon 2-6 deletion mice. **(C)** Western blot of P0 mouse tissue showing loss of Ret (red bands) expression in the *Ret* exon 2-6 deletion mice. *Gapdh* was used as a loading control (green bands).

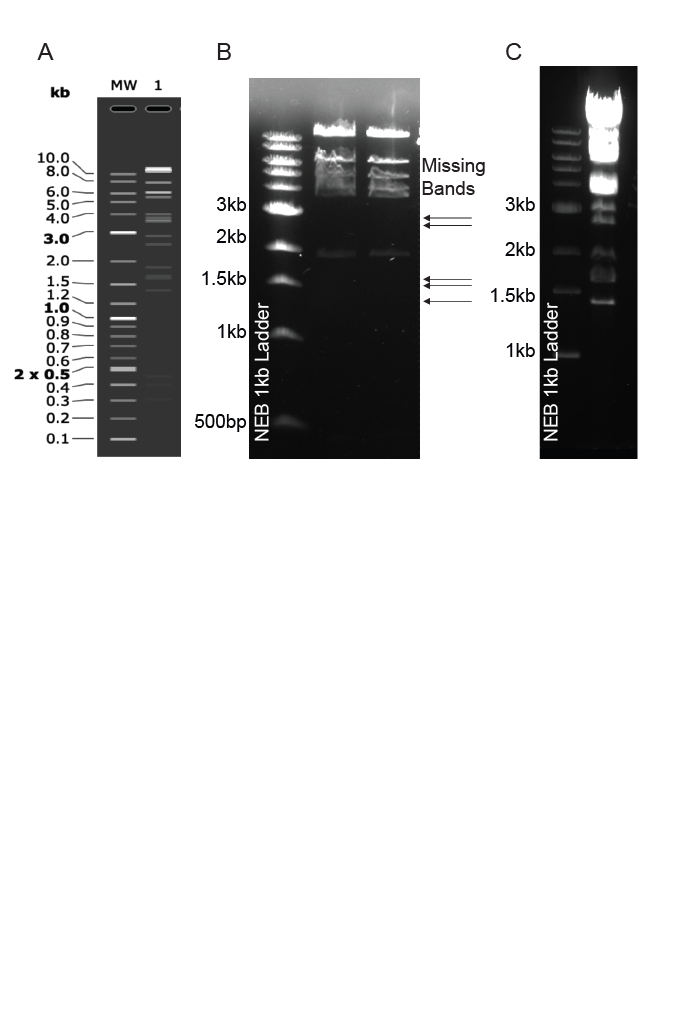

**Figure S4. Using BamHI digestion to distinguish bad and good pRDF90 delivery clones. (A)** *in silico* digestion of *RET* delivery BAC. **(B)** Example BamHI digestion pattern of two partially complete *RET* delivery BAC clones. **(C)** Example digestion of a complete *RET* delivery BAC clone.
